# A helical assembly of human ESCRT-I scaffolds reverse-topology membrane scission

**DOI:** 10.1101/2020.01.31.927327

**Authors:** Thomas G. Flower, Yoshinori Takahashi, Arpa Hudait, Kevin Rose, Nicholas Tjahjono, Alexander Pak, Adam L. Yokom, Xinwen Liang, Hong-Gang Wang, Fadila Bouamr, Gregory A. Voth, James H. Hurley

## Abstract

The ESCRT complexes drive membrane scission in HIV-1 release, autophagosome closure, MVB biogenesis, cytokinesis, and other cell processes. ESCRT-I is the most upstream complex and bridges the system to HIV-1 Gag in virus release. The crystal structure of the headpiece of human ESCRT-I comprising TSG101:VPS28:VPS37B:MVB12A was determined, revealing an ESCRT-I helical assembly with a 12 molecule repeat. Electron microscopy confirmed that ESCRT-I subcomplexes form helical filaments in solution. Mutation of VPS28 helical interface residues blocks filament formation *in vitro* and autophagosome closure and HIV-1 release in human cells. Coarse grained simulations of ESCRT assembly at HIV-1 budding sites suggest that formation of a 12-membered ring of ESCRT-I molecules is a geometry-dependent checkpoint during late stages of Gag assembly and HIV-1 budding, and templates ESCRT-III assembly for membrane scission. These data show that ESCRT-I is not merely a bridging adaptor, but has an essential scaffolding and mechanical role in its own right.

---

The Endosomal Sorting Complexes Required for Transport (ESCRTs) catalyze so-called “reverse topology” membrane scission events, defined as those in which budding is directed away from the cytosol ^1, 2^. Such events include, but are not limited to, multivesicular body (MVB) biogenesis, HIV-1 particle egress^3, 4^; cytokinesis ^5^; autophagosome closure ^6–8^; and nuclear envelope (NE) reformation^9, 10^. The core ESCRT machinery consists of the ESCRT-I, ESCRT-II and ESCRT-III complexes, ALIX, and the AAA^+^ ATPase VPS4. ESCRT-III and VPS4 are considered to be the minimal machinery for membrane scission, in that they can carry this reaction out *in vitro* in the absence of the upstream components ^11^. The ESCRT-IIIs are a family of small alpha helical proteins that can self-assemble into membrane-associated, homo- and hetero-polymeric structures. These polymeric assemblies can adopt many different morphologies, including cones ^12^, coils ^13^, flat spirals ^13–17^, and helical tubes ^12, 14, 18, 19^. The subunits within ESCRT-III polymers are exchanged with cytosolic monomers by VPS4^20^, leading to scission ^11^.

Viral proteins and host cargoes are recognized by the upstream ESCRT-I complex ^21–24^ or ALIX ^25, 26^. ESCRT-I in turn recruits ESCRT-II and then CHMP6, the most upstream of the ESCRT-III proteins. ALIX, and its paralog HD-PTP, provide a parallel route for ESCRT-III recruitment via CHMP4 ^27–30^. It is still unclear how ESCRT-I/-II and ALIX orchestrate ESCRT-III polymerization for scission. Each of these complexes are rigid and elongated, and their physical dimensions approximate those of the membrane necks with which they function. A central question is whether ESCRT-I (and other upstream ESCRTs) merely bridges cargo and the downstream ESCRTs, or whether it has a more specific and active mechanical role in scaffolding membrane remodeling and scission.

The critical Pro-Thr/Ser-Ala-Pro (PTAP) motif of HIV-1 Gag is recognized by the ubiquitin E2 variant (UEV) domain of TSG101 ^31–34^. Human ESCRT-I is a heterotetrameric complex consisting of a 1:1:1:1 complex of (a) TSG101, (b) VPS28, (c) one of VPS37A, B, C, or D, and (d) one of the MVB12A, MVB12B, UBAP1, UBA1L, or UMAD1^35–40^. The structure of the human core has not been determined, but TSG101, VPS28, and VPS37A-D have enough sequence homology to yeast Vps23, Vps28, and Vps37, respectively, to infer that similar general principles apply.

The crystal structure of the yeast ESCRT-I complex consisting of Vps23, Vps28, Vps37, and Mvb12 has an 18 nm rod-like core ^35^. With the inclusion of linkers and non-core domains, the yeast assembly spans up to 22 nm ^41^. The UEV domain is connected by a ∼90 residue Pro-rich linker to the stalk, which is followed by the fan shaped headpiece ^35, 42, 43^. The headpiece and stalk are collectively referred to as the core. The headpiece is capped by VPS28, which has a C-terminal alpha-helical bundle (VPS-CTD) that binds directly to the ESCRT-II complex and is distal to the UEV cargo recognition end ^35, 42, 44–46^. Yet, despite the similar function and nomenclature, human MVB12A/MVB12B and the other UBAP1-MVB12A associated (UMA) domain proteins ^37^ have no sequence homology to yeast Mvb12. They are integrated into ESCRT-I via their UMA domain ^39, 47^. Thus, with no reported structures including UMA sequences, nothing is known at the structural level about how the UMA-containing subunits are incorporated into ESCRT-I. This has left a major gap in understanding how human ESCRT-I complexes are organized.

Here we describe the structure of the human ESCRT-I headpiece including VPS37B and MVB12A. The structure reveals that human MVB12A and yeast Mvb12 bind to equivalent regions of the ESCRT-I head, albeit with altered conformations, and that MVB12A occludes the TSG101 PTAP motif. Upon determining this structure, we made the unexpected observation that the ESCRT-I head self-assembles into helical tubes within the crystal. This has led us to propose a model wherein cargo-dependent clustering of ESCRT-I triggers helix formation providing a nucleating platform upon which the downstream ESCRTs can assemble into membrane-deforming polymers. By mutating filament contact residues, we were able to test this model in HIV-1 release and autophagosome closure. These observations show that ESCRT-I is not merely an adaptor, but forms highly organized assemblies that template the initiation of membrane remodeling.

## Results

### Structure of the MVB12A-containing human ESCRT-I headpiece

Sequence alignments of metazoan MVB12 homologs showed that their UMA contains two distinct segments of conservation in its N- and C-terminal regions, respectively (Fig. 1a). We found that a 22-amino acid fragment of MVB12A (206-228; Fig. 1b) was sufficient to pull down the entire human ESCRT-I headpiece (Fig. 1c). This region is the most N-terminal of the two conserved segments and will subsequently be referred to as ‘UMA-N’. On the basis of this information, we co-expressed MVB12A UMA-N in complex with the trimeric human ESCRT-I (TSG101:VPS28:VPS37B) headpiece. We determined the crystal structure of the resulting ESCRT-I tetramer at 2.2 Å resolution using a search model derived from the Vps23:Vps28:Vps37 headpiece portion of the yeast ESCRT-I core (Fig. 1d). As expected, these three subunits (TSG101, VPS37B, VPS28) are similar to the yeast structure. The conformation of the novel component, MVB12A, resembles an ‘S’ shape comprised of two short 3_10_ helices followed by an anti-parallel beta-sheet formed with TSG101 (Fig. 1d-g). While yeast Mvb12 binds to the same region, it adopts an α-helical conformation (Fig. 1h)^35^. Like human MVB12A, this is followed by an antiparallel beta-sheet formed with Vps23, the yeast TSG101 ortholog.

**Fig. 1:**
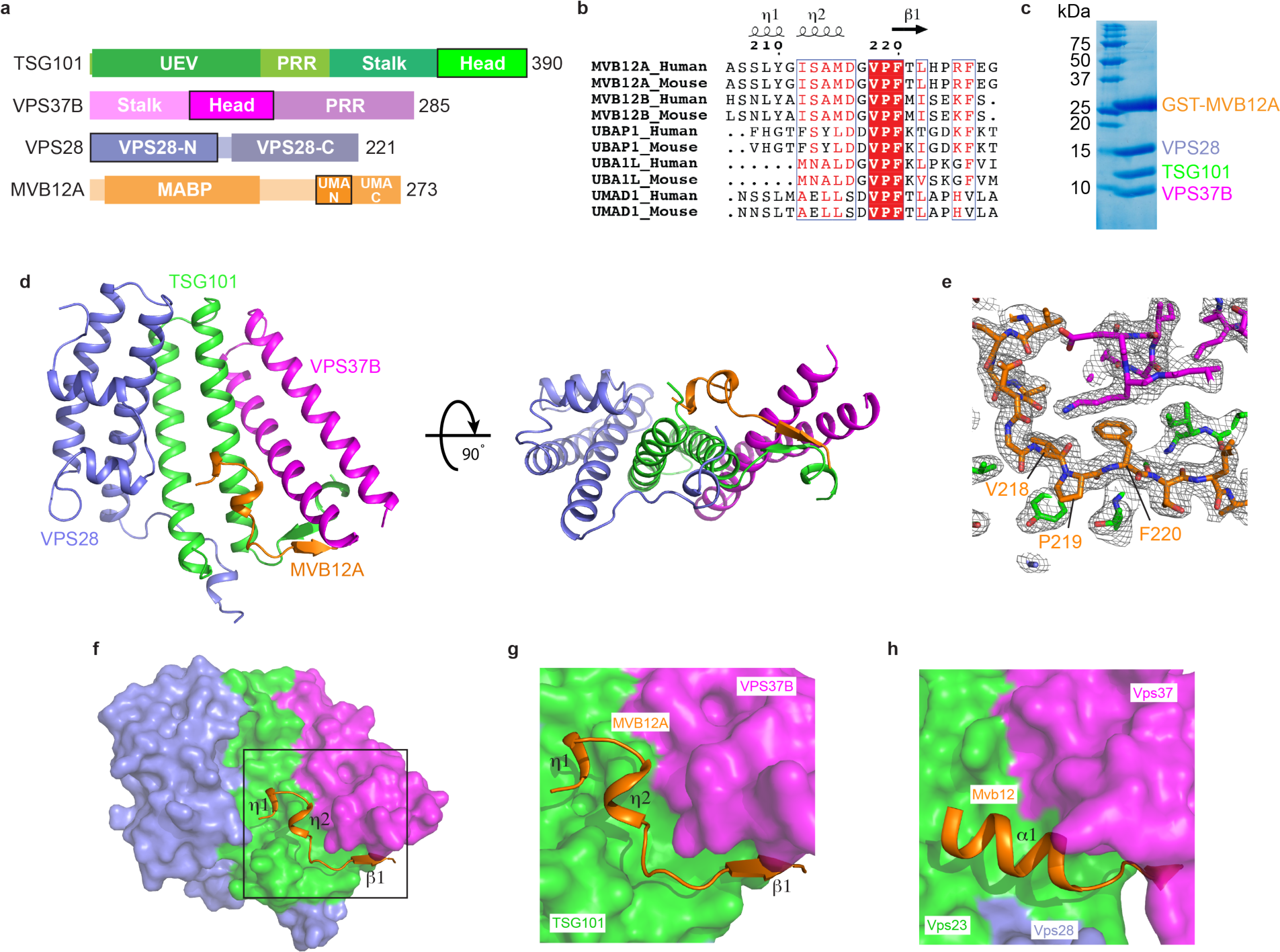
Structure of the heterotetrameric human ESCRT-I head a, Domain architecture of human ESCRT-I subunits. Constructs used for structural analysis are highlighted. **b,** Sequence alignment of MVB12A paralogs and orthologs. Secondary structure displayed above the alignment is derived from the human ESCRT-I head structure. Alignment was generated using ClustalW and ESPript. **c,** GST-tagged MVB12A 206-228 was co-expressed with trimeric ESCRT-I head (TSG101 308-388, VPS37B 97-167, VPS28 1-122) in *E.coli*. Lysate was incubated with glutathione beads and complex integrity analyzed by SDS-PAGE following multiple washes. **d,** Ribbon diagram of the human ESCRT-I head crystal structure. **e,** MVB12A 2Fo-Fc omit map. MVB12A chain was removed and model re-refined. Resulting 2Fo-Fc map is contoured at 1.0 σ and represented as a grey mesh. MVB12A chain replaced for clarity. **f,** Surface representation of the human ESCRT-I head with MVB12A shown as an orange ribbon. **g,** Enlargement of boxed region shown in (**f**). **h,** Mvb12 binding site from the previously determined heterotetrameric yeast ESCRT-I head (PDBID: 2P22). Mvb12, Vps37, Vps23 and Vps28 are colored orange, magenta, green and purple respectively.

Thus the mammalian UMA-N domain allows UMA-containing subunits to incorporate into the same site on the ESCRT-I headpiece as yeast Mvb12 despite the lack of any sequence homology and the differences in conformation.

### MVB12A interaction network in the headpiece

Within the UMA-N region, the residues VPF comprise a completely conserved bloc (Fig. 1b). The side chains of Val218 and Phe220 of the VPF motif protrude into a deep hydrophobic pocket formed by both TSG101 and VPS37B (Fig. 2a-c). The phenyl group of MVB12A Phe220 is wedged between Pro320 of TSG101 and the aliphatic chains of Lys155 and Lys158 and the side-chain of Val154 from VPS37B. MVB12A Val218 is buried against TSG101 Tyr325, Ile328, and Leu386. Pro219 is held in place by a hydrophobic interaction formed with Tyr325 of TSG101.

**Fig. 2:**
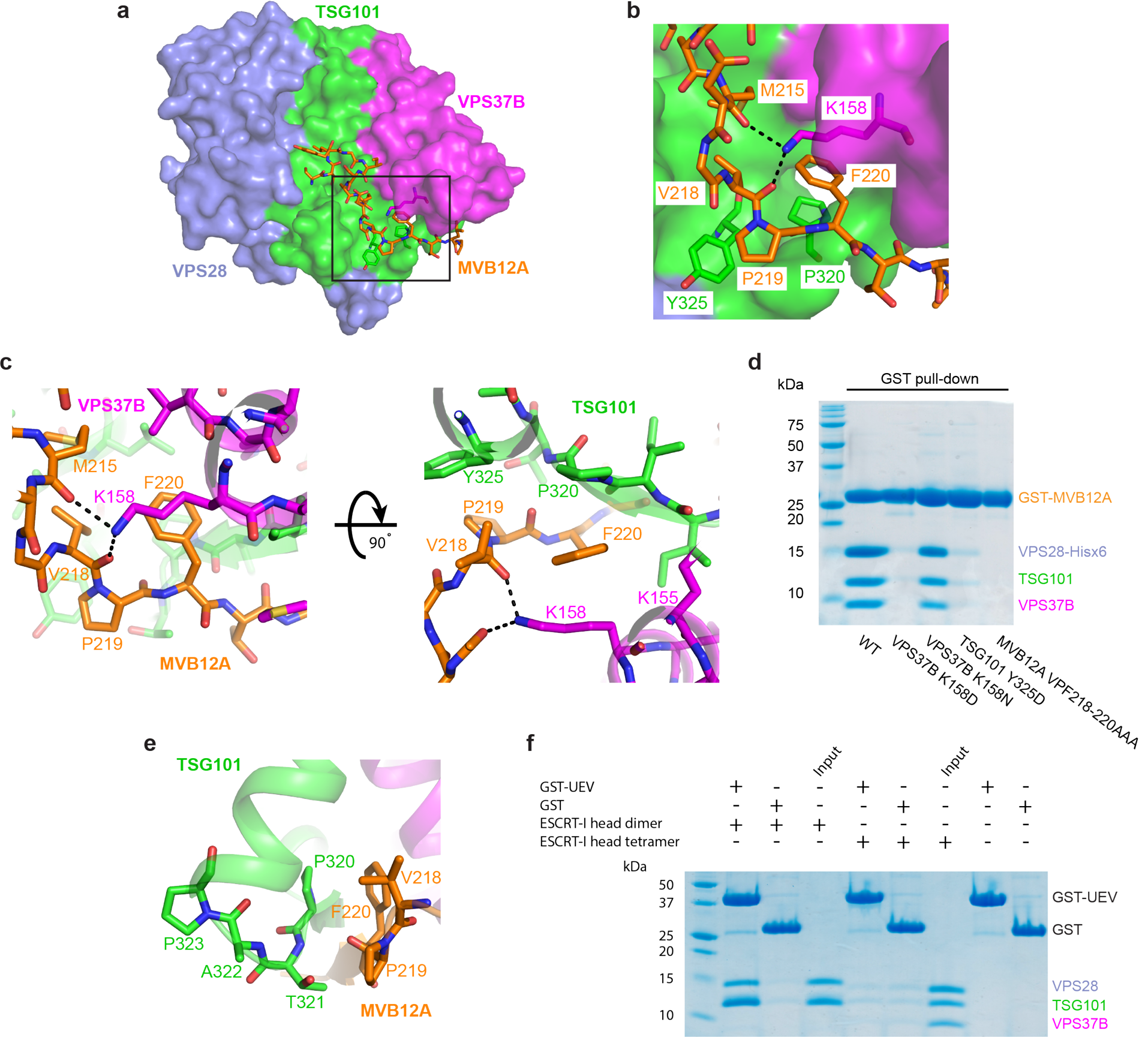
MVB12A VPF motif is required for the interaction with the ESCRT-I head and occludes the TSG101 PTAP motif **a,** Surface representation of the human ESCRT-I head with MVB12A and other key residues shown as sticks. **b,** Enlargement of boxed region in (**a**) showing the conserved MVB12A VPF motif binding pocket. TSG101, VPS37B, MVB12A colored green, magenta and orange respectively. Selected hydrogen bonds are shown as dashed lines. **c,** Stick representation of the MVB12A VPF motif binding pocket. **d,** Mutant versions of the GST-tagged ESCRT-I head complex were expressed in *E. coli.* Lysate was incubated with glutathione beads and complex integrity analyzed by SDS-PAGE following multiple washes. **e,** Interface between TSG101 PTAP motif and MVB12A VPF motif. **f,** GSH beads were coated with GST-TSG101 UEV or GST bait and incubated with binary (TSG101, VPS28) or tetrameric (TSG101, VPS28, VPS37B, MVB12A) ESCRT-I head. Beads were washed and complex formation analyzed by SDS-PAGE.

The side-chain of Lys158 donates multiple hydrogen bonds with the carbonyl groups of residues Met215 and Val218 of the MVB12A main-chain, helping to locking this region of main-chain in place. The contributions of these key residues to MVB12A integration into the headpiece were validated by in vitro pull-downs. Mutation of both TSG101 Tyr325 and VPS37B Lys158 to Asp essentially abolished the interaction with GST-MVB12A as judged by pull down experiments (Fig. 2d and Supplementary Fig. 1). Likewise, mutation of the VPF sequence to AAA also blocked incorporation (Fig. 2d and Supplementary Fig. 1).

Vertebrate TSG101 orthologs contain an internal PTAP motif that resides in the ESCRT-I headpiece. The UEV domain is capable of binding to peptides corresponding to the isolated TSG101 PTAP motif^48^. The human ESCRT-I headpiece structure revealed that access to the PTAP sequence of TSG101 is occluded by the VPF motif of MVB12A (Fig. 2e). The first Pro and the Thr of the PTAP motif form β-sheet hydrogen bonds with the Pro and Phe of the MVB12A VPF motif. These internal interactions made it seem unlikely that the UEV domain could bind to the PTAP motif in the context of this structure. In order to test this possibility, *in vitro* pull-down experiments were carried out using GST-tagged UEV as bait and various ESCRT-I head assemblies as prey (Fig. 2f). The UEV domain was capable of pulling down a dimeric version of the ESCRT-I head containing TSG101 and VPS28. This suggests that the PTAP motif of TSG101 is accessible to the UEV domain in the absence of MVB12A and VPS37B. Consistent with the observed sequestration of the PTAP motif in the full headpiece structure, the tetrameric version of the ESCRT-I head cannot bind to the UEV (Fig. 2e, 2f).

### Helical assemblies of ESCRT-I substructures in crystals and in vitro

Human ESCRT-I headpiece crystals grow in space group *P*6_1_ and contain six copies of the complex per asymmetric unit (Fig. 3a). This unusual combination of crystallographic and non-crystallographic symmetry elements corresponds to a crystal consisting of parallel helical tubes (Fig. 3b). Each tube is constructed of a series of three interwoven right-handed helices (Fig. 3b). The helices are built from individual ESCRT-I headpiece protomers that encircle a central lumen devoid of electron density. Each continuous helix consists of 12 ESCRT-I headpiece units per turn. The outer and lumen diameters of the assembly are 17 nm and 6.5 nm, respectively.

**Fig. 3:**
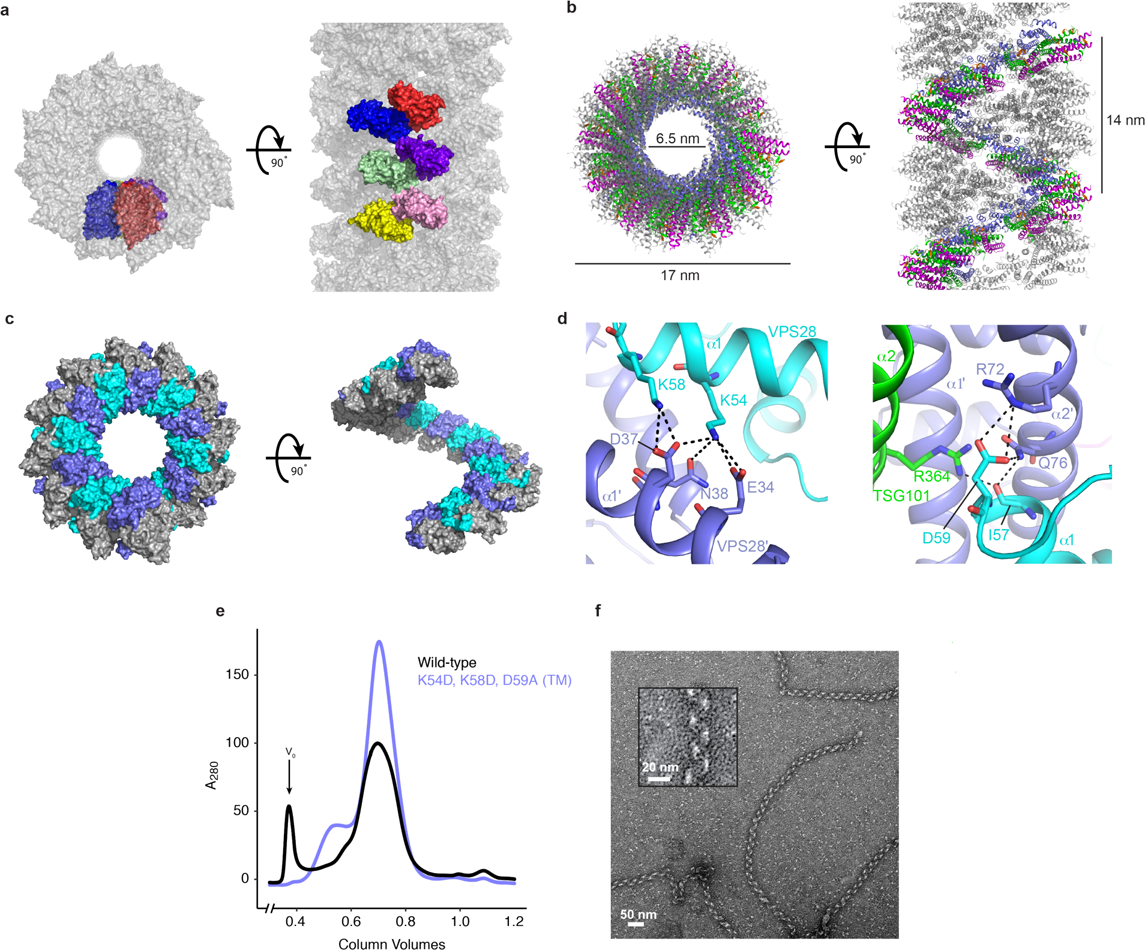
Helical ESCRT-I head assemblies **a,** The crystal from which the ESCRT-I head structure was determined consists of a series of helical tubes, one of which has been isolated and shown as a surface representation. The crystal exhibits *P*6_1_ symmetry with 6 copies of the tetrameric ESCRT-I head in the asymmetric unit. An individual asymmetric unit is highlighted and each ESCRT-I head protomer assigned a distinct color for clarity. **b,** Tube composed of three interleaved ESCRT-I head helices. One continuous helix is colored for clarity. TSG101, VPS28, VPS37B, MVB12A are colored green, purple, magenta and orange respectively. Tube dimensions are labelled. **c,** Surface representation of a single, continuous ESCRT-I helix. The intra-helical interface is primarily composed of residues from VPS28. Individual VPS28 subunits are colored either purple or cyan in order to clearly distinguish surface boundaries. **d,** Detailed views of the helical interface highlighting key residues. Inter-protomer bonds depicted with dashed lines. Subunits are colored as in (**c**). **e,** Size-exclusion chromatograms of WT and mutant (VPS28 K54D, K58D, D59A) binary (TSG101, VPS28) ESCRT-I head subcomplexes. Proteins were expressed in *E.coli* and subjected to initial IMAC purification prior to being applied on a size-exclusion column. Void volume (V_0_) labelled accordingly. **f,** Electron micrograph of WT ESCRT-I filaments obtained from the pooled V_0_ fraction shown in (**e**). No such assemblies were observed for the mutant construct. Zoomed view of a typical WT ESCRT-I filament shown inset.

Analysis of the intra-helical interface shows that it is mainly electrostatic in nature and is composed almost exclusively of residues from VPS28 (Fig. 3c and d). Most of these residues are well conserved from yeast to humans (Supplementary Fig. 2). VPS28 Lys54 and Lys58, located approximately midway along helix α1 of VPS28, form multiple salt bridges with acidic residues from the equivalent helix of a distinct copy of VPS28 (VPS28’) (Fig. 3d). Asp59, situated within a flexible loop linking α1 and α2 of VPS28, also forms multiple electrostatic interactions with Arg72 and Gln76 of VPS28’ α2’.

We noticed that similar interactions occur upon re-examining one of the yeast headpiece structures (PDB: 2CAZ) (Supplementary Fig. 3)^43^. The yeast ESCRT-I crystal is constructed from a series of laterally stacked tubes, each of which is composed of ESCRT-I head protomers arranged in a continuous helix. These tubes exhibit similar dimensions to their human counterparts with an outer diameter of 16 nm, inner diameter of 7.5 nm and an identical number of subunits per turn (12). The inter-protomer interfaces are also remarkably similar, although it was not possible to compare specific side chain interactions at the lower resolution of this yeast headpiece structure. The yeast tubes seen in the crystal contain a single, one-start, helix of ESCRT-I protomers as opposed to the three-start assembly seen in the human ESCRT-I crystal. As a result, the yeast assembly has a reduced helical pitch of 5 nm as compared with 14 nm for the human.

Attempts to purify an ESCRT-I headpiece subcomplex containing TSG101 and VPS28 subunits yielded a high molecular weight species as determined by size-exclusion chromatography (Fig. 3e). Isolation of this fraction and examination by negative stain electron microscopy identifies this higher molecular weight species as long, helical and filamentous in structure (Fig. 3f). These filaments are flexible, which precluded precise calculation of helical parameters and high-resolution cryo-EM structure determination. Nevertheless, the ∼20 nm outer diameter of these filaments is consistent, within the uncertainty of the measurement, with that of the human ESCRT-I headpiece filament in the crystal (Fig. 3a-c). In order to confirm that these filaments are analogous to the assemblies observed in the crystal structure, we produced a triple mutant VPS28 construct (K54D, K58D, D59A) designed to disrupt the helical interface, subsequently referred to as ‘VPS28 TM’. These mutants do not affect the ability of VPS28 and TSG101 to heterodimerize as judged by SDS-PAGE. However, the mutant complex failed to produce any higher molecular weight species, suggesting that the helical interface had indeed been disrupted and was the same as in the crystal structure. These results confirm that, like the ESCRT-IIIs, ESCRT-I subunits are capable of forming helical filaments in solution, and that these filaments are formed by electrostatic interactions between VPS28 protomers.

### The helical interface of VPS28 stabilizes ESCRT-I oligomerization *in vivo*

To examine the importance of the helical interface in ESCRT-I assembly *in vivo*, we examined the role of VPS28-mediated ring assembly in ESCRT recruitment to autophagosomes. HA-tagged VPS28 TM and control wild-type VPS28 (VPS28 WT) were lentivirally transduced into U-2 OS human osteosarcoma cells in which endogenous VPS37A is replaced by GFP-VPS37A^49^. Consistent with *in vitro* data, exogenously-introduced VPS28 displaced nearly all endogenous VPS28 and successfully incorporated into the ESCRT-I complex regardless of the mutations (Fig. 4a). The mutations also did not interfere with the membrane targeting of the complex as judged by the colocalization with HA-VPS28 MT and GFP-VPS37A on LC3-labelled autophagic structures (Fig. 4b). However, in the mutant-expressing cells, stabilization of ESCRT assembly by depleting the ESCRT-III component CHMP2A failed to induce further accumulation of VPS37A (Fig. 4c) despite an increase of autophagic structures (Fig. 4b, c).

**Fig. 4:**
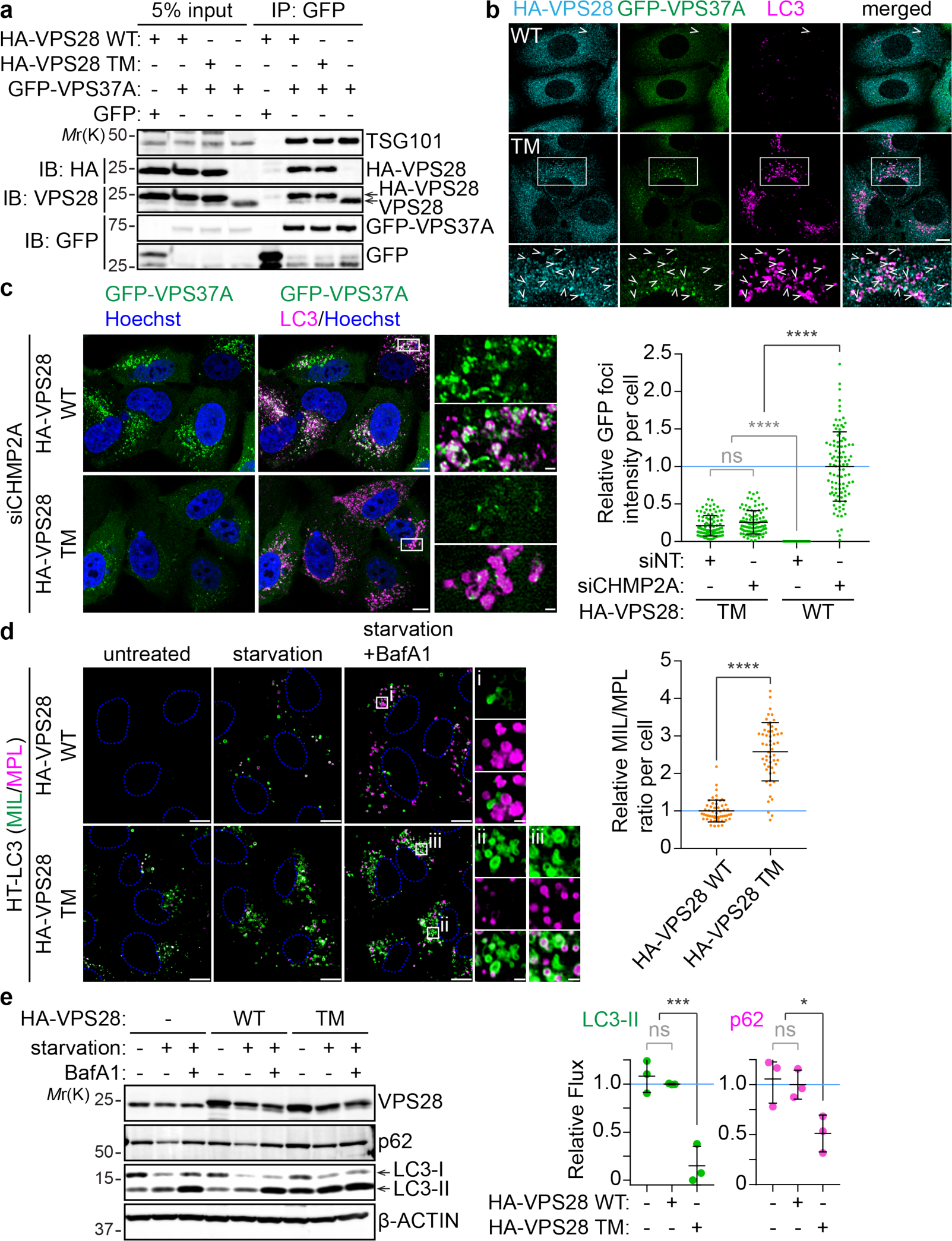
Helical contacts are required for phagophore closure in autophagy *VPS37A*-knockout (**a-c**) and wild-type (**d,e**) U-2 OS cells stably expressing GFP-VPS37A and HaloTag(HT)-LC3, respectively, were transduced with lentiviruses encoding HA-VPS28 WT or HA-VPS28 T-2 for 4 days. **a**, Cell lysates were immunoprecipitated with anti-GFP antibody and analyzed by immunoblotting using the indicated antibodies. **b**, Cells were starved for 3h, stained for HA and LC3, and analyzed by confocal microscopy. Arrowheads indicate colocalization of HA-tagged VPS28 (WT or T-2) with GFP-VPS37A on LC3-positive structures. **c**, Forty-eight hours after the transduction, cells were nucleofected with non-targeting siRNA (siNT) or CHMP2A siRNA (siCHMP2A) for 48 h, stained for LC3, and analyzed by confocal microscopy. Nuclei were counterstained with Hoechst 33342. The fluorescence intensity of GFP-VPS37A foci normalized to total GFP intensity per cell is shown relative to the mean of the control HA-VPS28 WT-expressing cells that were transfected with siCHMP2A (*n* = 98 cells). **d,e**, Cells were starved in the presence or absence of 100 nM Bafilomycin A1 (BafA1) for 3 h. **d**, Cells were subjected to the HT-LC3 autophagosome completion assay using AlexaFluor 488 (AF488)-conjugated membrane-impermeable ligand (MIL) and tetramethylrhodamine (TMR)-conjugated membrane-permeable ligand (MPL) and analyzed by confocal microscopy. The MIL/MPL fluorescence intensity ratio for each cell in the starvation plus BafA1 treatment group was calculated and shown relative to the mean of the control HA-VPS28 WT-expressing cells (*n* = 54 cells). **e**, Cell lysates were subjected to the immunoblotting-based autophagic flux assay using the indicated antibodies. Autophagic flux under starvation conditions was calculated as described in the *Methods* section and normalized to the mean of the control HA-VPS28 WT-expressing cells (*n* = 3 independent experiments). Representative blots or images from 2 (**a, b**) or 3 (**c-e**) independent experiments are shown. In **b**, and **c** and **d**, magnified images in the boxed areas are shown in the bottom panels and the right panels, respectively. The scale bars represent 10 µm, and 1 µm in the magnified images. Statistical significance was determined by one-way ANOVA followed by Holm-Sidak’s multiple comparison test (**c, e**) and Mann-Whitney nonparametric t-test (**d**). All values in the graphs are mean ± SD. ns, not significant; *, p≤0.05; ****, p≤0.0001.

These results indicate that the mutations within the helical interface of VPS28 disrupt the higher-order oligomerization of ESCRT-I without impairing the formation and membrane targeting of individual ESCRT-I tetramers.

### The ESCRT-I helical interface is required for autophagosome closure

The effect of ESCRT-I helical assembly disruption on phagophore closure was evaluated using the HaloTag-LC3 (HT-LC3) assay which employs the HaloTag receptor-conjugated LC3 reporter (HT-LC3) in combination with membrane-impermeable ligands (MIL) and -permeable ligands (MPL) labelled with two different fluorescent probes to distinguish membrane-unenclosed and -enclosed HT-LC3-II ^6^. To prevent the degradation of autophagosome-sequestered LC3-II, the experiment was performed in the presence of the lysosomal inhibitor Bafilomycin A1 (BafA1). As expected, the mutant-transduced cells displayed an accumulation of MIL^+^MPL^-^ unclosed autophagosomes and consequently increased the MIL to MPL ratio (Fig. 4d) indicating a defect in the membrane sealing. The importance of ESCRT-I helical assembly formation in autophagy was further determined by measuring autophagic flux. In the HA-VPS28 WT-transduced cells, starvation increased the amount of LC3-II which was further elevated in the presence of BafA1 similar to parental non-transduced control cells indicating normal autophagy induction and autophagic degradation (Fig. 4e). Consistently, the level of starvation-induced degradation of the autophagic substrate p62 in the WT-expressing cells was comparable to that in the control cells. In contrast, the levels of LC3-II and p62 were increased even under normal growth conditions and starvation-induced turnover of these proteins was significantly suppressed by the mutant expression. Collectively, these results indicate that ESCRT-I filament assembly is required for autophagosome closure and subsequent substrate degradation.

### The ESCRT-I helical interface is required for HIV-1 release

The role of the ESCRT-I helical assembly contacts in HIV-1 release was probed using the same triple mutant VPS28-TM described above. We used the previously described LYPxNL mutant (YP-) of the infectious pNL4.3 molecular clone, which lacks the Alix binding site and is therefore reliant only on the ESCRT-I, to promote virus release ^50^ to directly address the role of ESCRT-I in this process (Fig. 5). Endogenous VPS28 in 293T cells was first depleted by RNAi (Invitrogen) and after 48 h was replaced with either RNAi resistant WT VPS28 (lanes 3-5), an empty plasmid vector control (lane 2), or the TM VPS28 mutant (lanes 6-8) predicted to fail in forming higher order assemblies. As expected, reintroduction of WT but not mutant VPS28 led to a rescue in viral particle production by the YP-HIV-1 virus (lanes 3-5) similar to the non-depleted control (lane 1). Conversely, the VPS28 mutant supported very little to undetectable release of mature virus, indicating that the VPS28-mediated higher order assembly of ESCRT-I is essential for HIV-1 release.

**Fig. 5:**
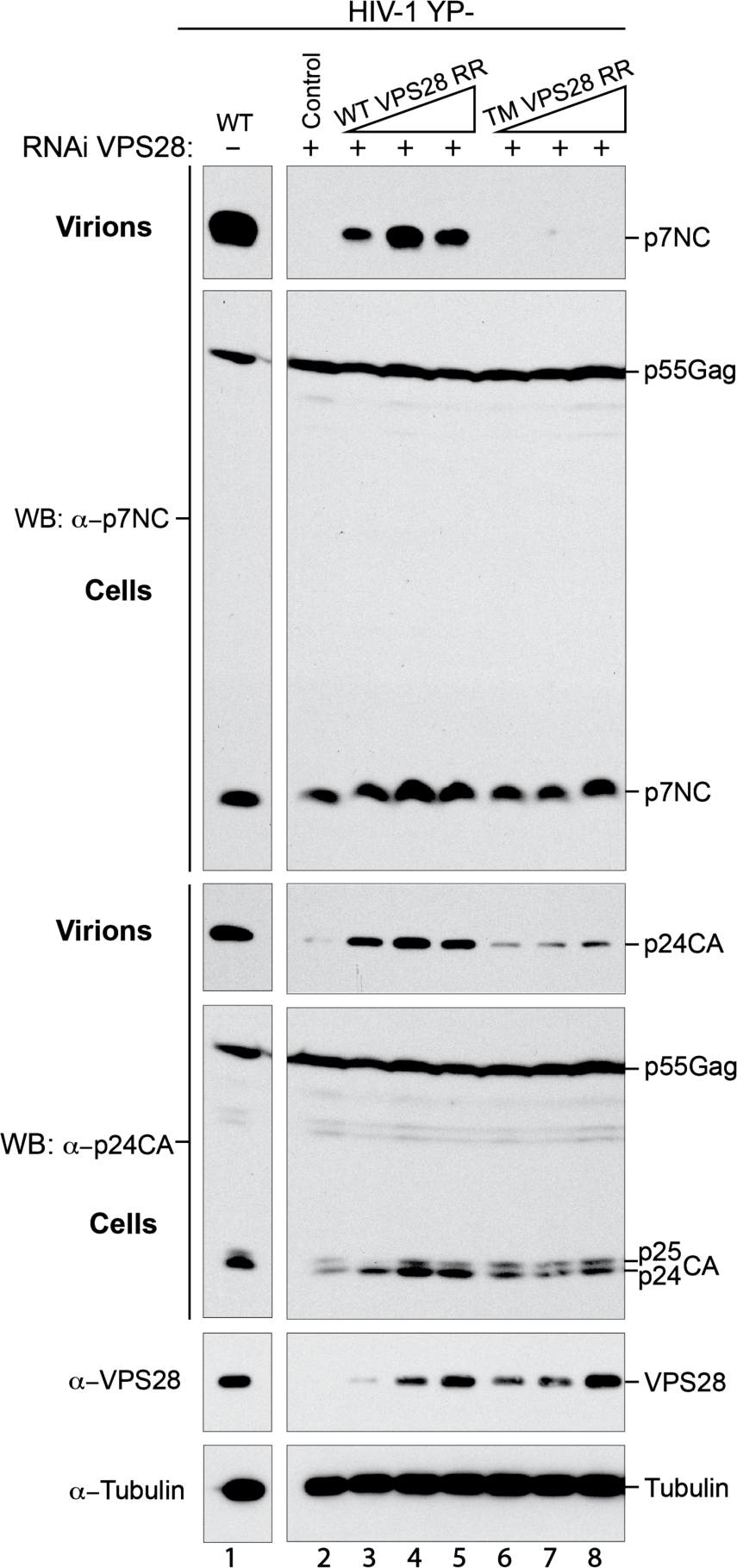
Helical contacts are required for efficient HIV-1 release Depletion of endogenous VPS28 with RNAi (lanes 2-8) and reconstitution with empty vector (lane 2), WT VPS28 (lanes 3-5), or VPS28 TM mutant (lanes 6-8) in 293T cells. Cells were co-transfected with an HIV-1 mutant with the LYPnXL L domain abrogated and virus release was compared with a non-depleted control (lane 1). Virion production and Gag expression levels were compared by Western blotting with anti-NC and anti-CA antibodies and cellular protein levels were probed with anti-VPS28 and anti-tubulin antibodies.

### Coarse-grained simulation reveals optimal Gag lattice geometry is essential for ESCRT-I oligomerization

To gain insight into the molecular structures of ESCRT-I assemblies at HIV-1 bud necks, we used coarse-grained (CG) molecular dynamics (MD) study to examine the formation of ESCRT-I rings at the necks of immature Gag lattice at various stages of initial budding. Our objective was to elucidate ESCRT-I assembly dynamics and determine the optimal Gag shell cross-section required for self-assembly of a 12-membered ESCRT-I ring oligomer. We first developed a CG model of ESCRT-I that is derived from the experimental structural data of the human ESCRT-I headpiece, yeast ESCRT-I stalk [PDBID: 2P22]^35^ and the ubiquitin E2 variant (UEV) domain of TSG101 [PDBID: 3OBU]^32^. The linker connecting the UEV domain of TSG101 and the stalk of ESCRT-I is modeled as a flexible linear polymer. The full-length ESCRT-I CG model used in this study is depicted in Fig. 6a. Our ESCRT-I CG model successfully captures the characteristic rod-like shape of the ESCRT-I core and the rotational flexibility of the ESCRT-I core imparted by the linker. We introduced attractive interactions between specific CG sites at the ESCRT-I head domain required for self-assembly. These CG sites correspond to amino acid residues of human ESCRT-I headpiece involved in direct electrostatic contacts in the helical assembly and predominantly belongs to VPS28. The location of energy minima corresponding to the attractive interactions between specific CG sites was set at the average distance between the same sites measured for the CG-representation of the 12 ESCRT-I headpiece units in the helical assembly. We chose the strength of these pairwise attractive interaction as *E*_E1-E1_ = −4 kcal mol^-1^. This particular value corresponds to the strongest interaction strength that does not lead to significant oligomerization in solution in our simulations (at least 90% of ESCRT-I hetero-tetramers do not oligomerize in solution, Fig. 6b). Finally, to emulate the recognition of the UEV domain of TSG101 by the PTAP motif of HIV-1 Gag, we included attractive interaction between the last bead of Gag and two CG sites at the C-terminal end of UEV domain. We describe the details of the ESCRT-I and Gag CG model in Methods.

**Fig. 6:**
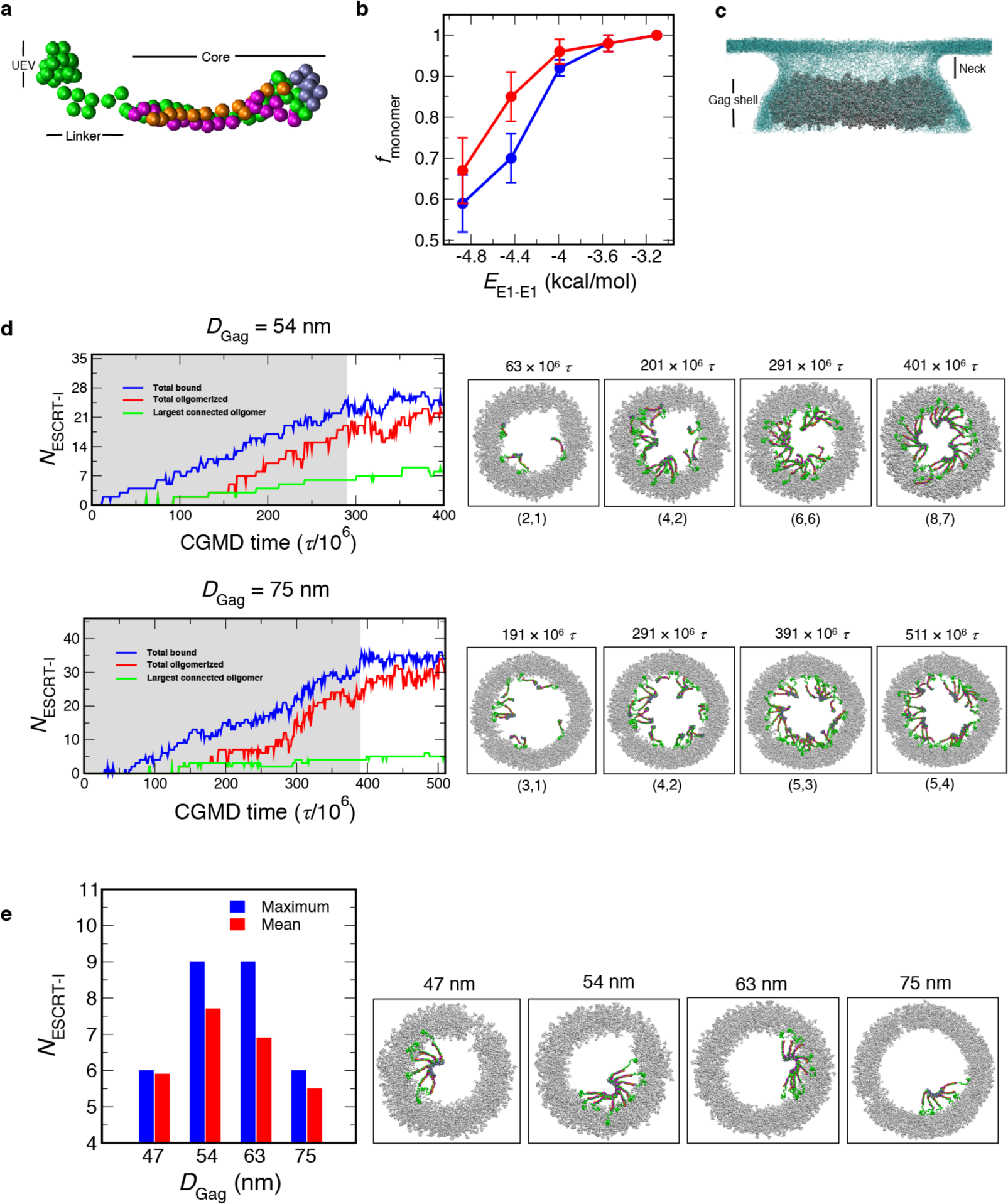
ESCRT-I assembly from CGMD simulations (**a**) CG representation of ESCRT-I. The CG sites representing TSG101, VPS28, VPS37B, MVB12A are colored in green, purple, magenta and orange respectively. (**b**) Weak interactions between ESCRT-I headpiece prevent oligomerization in solution. In a simulation cell containing 32 or 64 randomly placed ESCRT-I hetero-tetramers, we varied the strength of the ESCRT-I attractive interaction (*E*_E1-E1_) from −3.1 to −4.9 kcal/mol. The fraction of ESCRT-I hetero-tetramers remaining as monomer (*f*_monomer_) after 200 × 10^6^ *τ* monotonically decreases with increasing attractive interaction (*E*_E1-E1_). We note that *f*_monomer_ sharply decreases for *E*_E1-E1_ < −4 kcal/mol. The error bars represent standard deviation computed from the final outcome of 3 independent simulations. (**c**) Bud-like geometry used in the CGMD simulations. The Gag shell is represented in silver balls and the membrane is represented in light blue mesh. In our simulations ESCRT-I is introduced in the neck region. (**d**) ESCRT-I assembly time series profiles as a function of CGMD time step (*τ*) for Gag shells, *D*_Gag_ = 54 and 75 nm. The timeseries profile depicts (*i*) total ESCRT-I bound to Gag (blue), (*ii*) total oligomerized (red), and (*iii*) largest connected oligomer (green). The shaded region in the time series indicate insertion of ESCRT-I at the membrane neck every 10 × 10^6^ *τ*. The snapshots show a top view of ESCRT-I assembly at different stages of the time series. The size of the two largest oligomers are mentioned in parentheses for each snapshot. (**e**) Mean largest oligomer and observed maximum oligomer size for each Gag shell calculated from last 50 × 10^6^ *τ* of annealing CGMD trajectories. The snapshots depict the maximum largest oligomer observed for each Gag shell in the above-mentioned time frame.

To emulate the budding configuration of immature Gag lattice, we created model Gag shells of varying cross-sections representing immature Gag lattice at different stages of budding. The internal diameter (*D*_Gag_) of these Gag shells range from 47 nm to 75 nm. These Gag shells are wrapped with membrane resembling the budding neck geometry (Fig. 6c). The details of system preparation and equilibration is provided in Methods. To simulate the gradual accumulation of ESCRT-I to the site of Gag assembly we inserted a single ESCRT-I hetero-tetramer every 10 × 10^6^ CGMD time steps (*τ*) at the membrane neck region (Fig. 6c, Supplementary Movie 1). This particular frequency of ESCRT-I insertion was chosen to allow sufficient time for the inserted ESCRT-I to diffuse and reach the Gag shell. We inserted ESCRT-I hetero-tetramer until the lumen of the Gag shell is saturated and ESCRT-I is recruited by the Gag shell. These simulations ranged from 270 × 10^6^ to 420 × 10^6^ *τ* depending on the cross-section of the shell. The systems were further evolved for 120 × 10^6^ *τ* to allow for annealing of initial clusters formed during ESCRT-I recruitment and growth of the largest cluster. In what follows we summarize the assembly dynamics of ESCRT-I oligomerization.

To examine the assembly dynamics of ESCRT-I oligomerization we present *N*_ESCRT-I_ time evolution of the simulation trajectory of three parameters: (*i*) total ESCRT-I bound to Gag, (*ii*) total oligomerized, and (*iii*) largest connected oligomer. Fig. 6d shows the *N*_ESCRT-I_ time profile for a Gag shell of diameter (*D*_Gag_) 54 and 75 nm. In the initial phase of the simulation as ESCRT-I is introduced in the membrane neck we observe continuous recruitment of ESCRT-I by the Gag shell. In the course of the simulation and concurrent to ESCRT-I binding to Gag, ESCRT-I oligomerization progresses via attachment of bound monomers, as illustrated in Fig. 6e. We note that in the simulations, multiple clusters independently grew and persisted over the course of the simulations. These intermediate clusters are typically smaller than the 12-membered ring of ESCRT-I hetero-tetramers observed in experiments. Annealing to a complete 12-membered oligomer requires coalescence or partial disassembly of these intermediate clusters. In the second phase of the simulations and once lumen saturation was achieved, the systems were evolved without further insertion of any ESCRT-I hetero-tetramers. On the time scale of our CGMD simulations and for each Gag shell, we observed growth of the largest oligomer by rearrangement of the intermediate clusters. Overall, our simulations suggest a slow and stepwise pathway of ESCRT-I oligomerization. In what follows, we summarize the outcome of our assembly simulations for Gag shells of different cross-sections and report the average largest oligomer to illustrate the efficacy of ESCRT-I self-assembly for Gag shells of different diameters.

Fig. 6e shows the mean largest ESCRT-I oligomer size and the maximum oligomer size for each Gag shell of diameter *D*_Gag_ = 47, 54, 63 and 75 nm. For each Gag shell, the largest oligomer is calculated over the last 50 × 10^6^ *τ* of the annealing simulations and averaged over 3 independent simulations. On the time scale accessible to our simulations, we can observe the growth of ESCRT-I oligomer up to a maximum 9-mer for *D*_Gag_ = 54 nm. which is close to the experimental 12-mer ESCRT-I helical assembly. In contrast, Gag shells *D*_Gag_ = 63 and 75 nm exhibit diminished assembly competence – the wider the cross-section of the shell, the less the geometric compatibility with higher-order ESCRT-I oligomers. Similarly, ESCRT-I oligomerization is also hindered by a narrower Gag shell (*D*_Gag_ = 47 nm). Our simulations illustrate that an optimal Gag shell is required to stabilize higher-order ESCRT-I assembly. Based on our simulations, we expect that assembly of the full 12-mer ESCRT-I complex is templated when the opening of the immature Gag virion approaches 50 nm during budding. We note that our CGMD simulations likely cannot access the timescales required for extensive partial disassembly and assembly of intermediate clusters typical for annealing, thereby leading to the perfect 12-ESCRT-I oligomer.

## Discussion

The structure of the heterotetrameric human ESCRT-I headpiece sheds new light both on the organization of ESCRT-I itself, and on the larger question of how the ESCRT system as a whole orchestrates membrane scission. With respect to the first point, the structure answered the long-standing question as how UMA domain proteins are incorporated into the mammalian complex revealing that the UMA domain of MVB12A occupies what is essentially a cognate location in the headpiece as does yeast Mvb12. Yet the overall secondary structure of MVB12A is distinct, as are the interactions its makes with the headpiece. In the structure, the VPF motif of MVB12A UMA-N region is the main anchor. The VPF motif is conserved throughout the UMA domain proteins, thus this structure should serve as a prototype for the five different human UMA proteins and the twenty theoretically possible human ESCRT-I complex variants. While it has been reported that some of these theoretical complexes do not actually form ^39^, the VPF motif and the sites that it contacts on TSG101 on VPS37B are well conserved, and seem unlikely to be complex-specific determinants. Subcomplex specificity is therefore likely to be determined by residues in the stalk or elsewhere.

The UEV domain of TSG101 has been shown to bind to the TSG101 PTAP motif, which could in principle autoinhibit ESCRT-I with respect to cargo recognition^48^. However, interactions with the VPF motif sterically occlude the internal TSG101 PTAP motif. Our pull-down experiments confirmed that while the UEV domain of TSG101 is capable of binding to a dimeric version of the ESCRT-I headpiece consisting of just TSG101 and VPS28, it is unable to bind to the tetrameric complex. The subunits of most protein complexes are not synthesized with precisely matching stoichiometry, and partially assembled complexes can act as dominant negatives.

Exposure of the PTAP motif in the partially assembled TSG101:VPS28 complex could potentially serve to prevent premature cargo recognition by an immature precursor complex. Upon formation of the active tetrameric complex MVB12A would sequester the internal PTAP motif, dislodge the UEV domain, and so free it up for cargo recognition.

The VPF motif also interacts with Lys158, a residue that is highly conserved amongst metazoan VPS37B homologs. Homozygous mutation of the equivalent residue within VPS37A to asparagine has been linked with complex hereditary spastic paraparesis (CHSP) in multiple individuals^51^. It is interesting that VPS37A is the ortholog uniquely involved in autophagosome closure ^49^, but it is currently whether or how this role is connected to the disease phenotype. In our pull-down experiments the K158N mutation had little effect on the interaction between MVB12A UMA-N and the trimeric head while K158D abolished the interaction. Complete abolition of the UMA protein incorporation into VPS37A complexes by K158D would have a stronger defect, perhaps resulting in embryonic lethality as seen with TSG101^52^, as opposed to the CHSP resulting from K158N.

The ESCRTs are an integrated pathway consisting of the upstream complexes ESCRT-I and -II and ALIX, and the downstream membrane-severing ESCRT-III complex and the VPS4 ATPase. Throughout the many ESCRT-dependent cell processes, the upstream and downstream components work together to carry out biologically important membrane neck scission events. In one of the very few pathways that appear to be independent of upstream complexes, nuclear pore reformation, the key ESCRT-III subunit CHMP7 contains an ESCRT-II VPS25-related domain. Thus, even in nuclear envelope reformation, a structure related to an upstream complex is present. Despite that both upstream and downstream components are required for biological function, nearly all structural ^12^ and biophysical ^20, 53^ analysis of higher order assemblies of ESCRTs has focused on the downstream components. Structures have been observed in cellular cryo-ET ^54^ and deep etch EM (DEEM) ^17^ that resemble the dimension of a postulated bud neck assembly of ESCRT-I ^55^. However, the resolution of these studies has not been sufficient to positively identify ESCRT-I, nor to design mutational probes. Thus, our understanding of how ESCRT-I (and ESCRT-II and ALIX) might be arranged in three dimensions at ESCRT membrane scission sites has lagged behind that for ESCRT-III.

In the course of solving the crystal structure of the human ESCRT-I headpiece, we identified a higher-order, helical ESCRT-I assembly. The pitch of the helix is variable when comparing yeast and human structures. There are no direct contacts between successive turns of the human ESCRT-I helix, there it seems possible, or even likely that in cells, a single ring rather than a helix, is formed. A recent estimate of the number of copies of TSG101 at the HIV-1 bud neck is consistent with a single ring ^56^. Using this assembly as a template upon which to model the full complex, the UEV domains project outward to form a 12-membered ring. This dimension is plausible for many of the cell processes involving ESCRT-I, including MVB budding, autophagosome closure, and HIV-1 release. In HIV-1 release, this dimension is about half of the maximum inner diameter of the immature Gag lattice. Thus it corresponds to an immature lattice has grown past a hemisphere and from there, half-way to complete closure. Electron-microscopy-based imaging studies have identified spoke-like projections of approximately 20 nm in length that appear to localize at the edge of the immature Gag lattice ^17, 54^. These assemblies are only observed in cells when the bud neck is more than half the diameter of the bud, suggesting that they assemble during the early stages of bud formation ^54^. Similarly, in the coarse grained simulations reported here, larger assemblies of ESCRTs are only stable when the diameter of the bud neck is close to the most preferred value. If the ESCRTs were to act at this stage, this would leave a corresponding gap in the immature lattice, consistent with observation^57^. This would also result in incorporation of a small number of ESCRT-I complexes into the released virion, consistent with reports ^56, 58^.

The VPS28 C-terminal domain (CTD) responsible for recruiting downstream factors is connected to the headpiece portion of VPS28 by an 11 amino acid linker. Given the short linker and the small mass of the CTD (11.5 kDa), the diameter of the complete ESCRT-I ring including the CTD should be very close to what is seen here with the headpiece alone. With inner and outer diameters of 6.5 and 17 nm, respectively, the headpiece ring dimensions correspond to the narrow ends of ESCRT-III cone structures that have been published ^12, 59^. Most tubular ESCRT-III structures are larger, if not much larger, with diameters of 24 nm to 400 nm ^12, 60^. The narrowness of the ESCRT-I templating ring suggests that the ESCRT-III cone grows outward from the ESCRT-I ring, rather than inward ^1^, ^2, 61^. This might be consistent with the return of most of the ESCRT-III to the cytosol, but with the ESCRT-I retained in the virion ^2–4^. The same applies to physiological functions such as autophagosome closure.

This leads us to propose the following model for the role of ESCRT-I in HIV-1 release. The general outlines would apply to autophagosome closure and MVB budding as well, although it is still unknown how ESCRT-I is targeted to autophagosomes. Prior to HIV scission, single ESCRT-I tetrameric complexes are gradually recruited to Gag ^62^. In analogy to this, we observed that VPS28 mutations do not affect recruitment of ESCRTs to LC3 membranes. Without a geometry compatible with 12-membered ring formation, they do not template ESCRT-III spirals, consistent with the observation that ESCRT-III is not stably recruited at this stage ^63^ despite the presence of ESCRT-I. As Gag budding proceeds and the cross-section of the membrane neck narrows to around 55 nm, ESCRT-I at the Gag boundary reaches high local concentration and the geometry is compatible with ring formation, stabilizing the assembly and preventing dissociation. Indeed, HIV budding profiles with openings of about these dimensions have been reported to manifest spoke-like assemblies ^54^. This leads to the creation of a helical or ring-shaped platform, upon which the downstream ESCRTs assemble to drive membrane scission with the help of VPS4. This nucleation model could explain why live cell imaging studies have reported a gradual accumulation of ESCRT-I at budding sites ^62^ followed by a sharp late burst in recruitment of the ESCRT-IIIs immediately prior to scission ^63^. It also explains why CHMP2A depletion does not lead to upstream ESCRT accumulation in VPS28 mutant cells in our experiments. Our finding that the filament contacts are conserved from yeast to humans, are functionally required for autophagosome closure, and explain ESCRT recruitment patterns in both HIV release and autophagosome biogenesis, suggests this mechanism applies widely.

## Online methods

### Protein expression and purification

The codon-optimized synthetic genes for human TSG101 308-388, VPS37B 97-167, VPS28 1-122 and MVB12A 206-268 were subcloned into cassettes 1-4 of the polycistronic pST39 vector. MVB12A was tagged with an N-terminal GST tag followed by a TEV cleavage site. The plasmid was transformed into *E. coli* strain BL21(DE3) Star followed by overnight expression at 20 °C. Cells were pelleted and resuspended in buffer (50 mM Tris pH 8.0, 300 mM NaCl, 3 mM β-mercaptoethanol, 1 mM phenylmethylsulfonyl fluoride) and lysed by sonication. The tetrameric complex was isolated by glutathione affinity chromatography followed by overnight on-resin TEV cleavage. Cleaved product was dialyzed into low salt buffer (50 mM Tris pH 8.0, 50 mM NaCl, 3 mM β-mercaptoethanol), applied to a HiTrap Q-sepharose FF column (GE Healthcare) and eluted with a linear 50 mM to 1 M NaCl gradient. Pooled fractions were passed through nickel-charged agarose to remove residual hexahistidine-tagged TEV protease. Flow-through was concentrated and applied to a Superdex 75 size exclusion column. The fractions containing the complex were concentrated in buffer (20 mM Tris-HCl pH 8.0, 150 mM NaCl, 0.5 mM TCEP).

The binary TSG101, VPS28 subcomplex was cloned in a similar manner to the tetrameric complex but VPS37B and MVB12A were omitted and a TEV-cleavable hexahistidine tag was fused to the C-terminus of VPS28. The subcomplex was expressed in BL21(DE3) *E. coli* cells Star at 20 °C overnight. Cells were resuspended in buffer (50 mM Tris-HCl pH 8.0, 300 mM NaCl, 3 mM β-mercaptoethanol, 1 mM phenylmethylsulfonyl fluoride, 10 mM imidazole) and lysed by sonication. The ESCRT-I head complex was isolated by Ni^2+^ affinity chromatography, washed with 30 mM imidazole buffer and eluted with 300 mM imidazole. Eluant was concentrated and applied to a Superdex S200 size exclusion column equilibrated in buffer (20 mM Tris-HCl pH 8.0, 150 mM NaCl, 0.5 mM TCEP). The subcomplex eluted as a two peaks, each of which was individually pooled.

DNA coding for the human TSG101 UEV domain (residues 2-145) was subcloned into the pGST2 vector. The plasmid was transformed into E. coli strain BL21(DE3) Star and expressed overnight at 30 °C. Cells were pelleted and resuspended in buffer (50 mM Tris pH 8.0, 300 mM NaCl, 3 mM β-mercaptoethanol, 1 mM phenylmethylsulfonyl fluoride) and lysed by sonication. GST-UEV was isolated using glutathione affinity chromatography, washed, and eluted with buffer containing 30 mM reduced glutathione. Eluate was concentrated and further purified using a Superdex 200 size exclusion column equilibrated in buffer (20 mM Tris pH 8.0, 150 mM NaCl, 0.5 mM TCEP).

All mutant constructs used in this study were generated by QuikChange site-directed mutagenesis according to the manufacturers instructions (Agilent).

### X-ray crystallography

Crystals of human ESCRT-I head were grown using the hanging drop vapor diffusion method at 18 °C. 2 µL of the protein sample (8.2 mg ml^-1^) was mixed with 2 µL reservoir solution and suspended over a 500 µL reservoir of 2 % w/v γ-Polyglutamic acid (γ-PGA) (Molecular Dimensions), 100 mM sodium formate, 100 mM sodium acetate pH 5.0. Crystals appeared within 3 days and continued to grow for approximately one week. Crystals were cryoprotected in reservoir solution supplemented with 25 % (v/v) Glycerol. A Native dataset was collected from a single crystal under cryogenic conditions (100 K) at a wavelength of 1.11583 Å using a Dectris Pilatus3 S 6M detector (beamline 8.3.1, ALS). The data was indexed and integrated using XDS^64^. Integrated reflections were scaled, merged and truncated using the CCP4 software suite ^65^.

Initial phases were determined by molecular replacement with the program PHASER^66^ using the trimeric yeast ESCRT-I head structure (PDBID: 2F66) as a search model. Iterative rounds of manual model building and refinement were performed using Coot^67^ and Phenix Refine^68^ respectively. 99.87% of the residues are in the most favored and additionally allowed regions of the Ramachandran plot. The space group is P6_1_ with 6 molecules per asymmetric unit. Statistics of the final structure are shown in (Supplementary Table 1). Structural figures were produced using the program PyMOL (W. Delano, https://pymol.org).

### Electron microscopy

The ESCRT-I TSG101, VPS28 head subcomplex was purified as described above. Void peak fractions from the final size-exclusion step were pooled and diluted to 3.5 µM for negative stain electron microscopy. 5 µl of sample was applied to glow-discharged continuous carbon-coated copper grids (CF400-CU, Electron Microscopy Sciences) and negatively stained with 2 % (w/v) uranyl acetate. Samples were inspected using transmission electron microscopy (Tecnai-12, FEI) operated at 120 keV with a magnification of 49,000×. Images were collected with a charge-coupled device (CCD) detector (4k TemCam-F416, TVIPS).

### Pull-down assays

Pull-downs were performed to test the effect of specific mutations on the ability of MVB12A to interact with the ESCRT-I head. In the case of both the WT and mutant constructs, GST-MVB12A (206-268) was co-expressed with TSG101 (308-388), VPS37B (97-167) and a C-terminal hexahistidine tagged version of VPS28 (1-122). Cell lysates were applied to either 30µL of GSH or Ni-NTA resin prewashed with binding buffer (20 mM Tris-HCl pH 8.0, 150 mM NaCl, 0.5 mM TCEP, 20 mM imidazole) and incubated at 4 °C for 30 minutes. The beads were washed three times and complex formation analyzed by SDS-PAGE.

For the TSG101 UEV pull-down assays, pre-purified bait (GST-UEV or free GST) was incubated with 30 µL of GSH resin prewashed in binding buffer (20 mM Tris-HCl pH 8.0, 150 mM NaCl, 0.5 mM TCEP) for 30 minutes at 4 °C. Beads were washed three times to remove excess bait. ESCRT-I head prey (tetramer or dimer) was added to the beads in 500 µL binding buffer and incubated at 4 °C overnight. Beads were again washed three times to remove unbound prey and bound proteins analyzed by SDS-PAGE.

### Antibodies and siRNAs

The following antibodies were used for immunoblotting (IB) and immunofluorescence (IF): β-ACTIN (IB, Sigma-Aldrich, A5441, 1:10,000); GFP (IB, Cell Signaling, 2956, 1:1,000); HA (IB, IF, BioLegend, 901513, 1:2,000 (IB), 1:1,000 (IF)); MAP1LC3B (IB, Novus, NB100-2220, 1:3,000; IF, Cell Signaling, 3868, 1:200); p62 (IB; American Research Products, 03-GP62-C, 1:4,000); TSG101 (IB, Abcam, ab83, 1:1,000); VPS28 (IB, Santa Cruz Biotechnology, sc-166537, 1:100). ON-TARGETplus SMART Pool Non-targeting (D-001810-10) and CHMP2A (L-020247-01) siRNAs were obatined from GE Healthcare Dharmacon. pCDH1-CMV-HA-VPS28(WT)-SV40-hygro and pCDH1-CMV-HA-VPS28(TM[K54D, K58D, D59A])-SV40-hygro were generated using Gibson Assembly. All other reagents were obtained from the following sources: Bafilomycin A1 (LC Laboratories, B-1080); Hoechst 33342 (Invitrogen, NucBlue, R37605); Membrane-impermeable HaloTag Ligand (MIL) (Promega, Alexa Fluor 488-conjugated, G1001); Membrane-permeable HaloTag Ligand (MPL) (Promega, tetramethylrhodamine-conjugated, G8251); normal goat serum (Sigma-Aldrich, G9023); Nucleofector Kit V (Lonza, VCA-1003); paraformaldehyde (Electron Microscopy Sciences, 15710); XF Plasma Membrane Permeabilizer (XF-PMP) (Seahorse Bioscience, 102504-100).

### Cell culture

U-2 OS cells were obtained from American Type Culture Collection and maintained in McCoy’s 5A Medium supplemented with 10% FBS. To induce autophagy, cells were rinsed three times with Dulbecco’s Phosphate Buffered Saline (DPBS) and incubated with amino acid-free DMEM (Wako, 048-33575). GFP-VPS37A-expressing *VPS37A* knockout U-2 OS cells ^6^ and HT-LC3-expressing U-2 OS cells ^49^ were generated as described previously.

### Coimmunoprecipitation

Cell lysates were prepared in 0.5% NP-40 lysis buffer (10 mM Tris/Cl pH 7.5, 150 mM NaCl, 0.5 mM EDTA, 0.5% NP-40) containing protease inhibitors and subjected to immunoprecipitation using with anti-GFP-conjugated agarose beads (GFP-Trap beads, Chromotek, gtma). The immunoprecipitates were washed three times with lysis buffer and subjected to immunoblotting.

### Immunofluorescence and confocal microscopy

Cells grown on Lab-TekII Chambered Coverglass, Chamber Slide (Nunc, 154941) were fixed and permeabilized in 4% paraformaldehyde-PBS at room temperature (RT) for 3 min followed by methanol at −20°C for 7 min, blocked in 10% normal goat serum for 1 h, and incubated with the primary antibodies overnight at 4°C followed by the secondary antibodies for 1h at RT. All fluorescence images were obtained using a Leica AOBS SP8 laser-scanning confocal microscope (63x oil-immersion [1.2 numerical aperture] lens) with the highly sensitive HyD detectors and the Leica image acquisition software LAX, deconvolved using Huygens deconvolution software (Scientific Volume Imaging), and analyzed using Imaris software (Bitplane) and Volocity software (PerkinElmer) without gamma adjustment.

### Autophagy assays

To induce autophagy, cells were rinsed three times with Dulbecco’s Phosphate Buffered Saline (DPBS) and incubated with amino acid-free DMEM (Wako, 048-33575) for 3 h. HT-LC3 autophagosome completion assay was performed as described previously ^49^. For immunoblotting-based autophagic flux assay, cell lysates were prepared in radio-immunoprecipitation assay buffer (150 mM NaCl, 10 mM Tris-HCl, pH 7.4, 0.1% SDS, 1% Triton X-100, 1% Deoxycholate, 5 mM EDTA, pH 8.0) containing protease inhibitors and subjected to SDS-PAGE followed by immunoblotting with antibodies against LC3, p62 and β-ACTIN. The signal intensities were quantified using the Image Studio version 5 software (LI-COR Biotechnology) and the levels of LC3-II and p62 were normalized to the respective value of β-actin. Autophagic flux was calculated by subtracting normalized LC3-II and p62 values in starvation group from those in starvation plus BafA1 group.

### Statistical analyses

Statistical significance was determined using Graph Pad Prism 7.0. Threshold for statistical significance for each test was set at 95% confidence (p<0.05).

### HIV-1 release

293T cells (2.2 × 10^6^) were seeded into T25 flasks and transfected with VPS28-specific RNAi or control RNAi (Invitrogen), 1-2 µg of LYPxNL-abrogated (YP-) mutant HIV-1 proviral DNA together with expression vectors for RNAi Resistant (RR) wild-type (WT) or mutant VPS28, (TM) or an empty vector control. Twenty-four hours after transfection, culture supernatants were harvested as previously described in ^69^ and virions were then pelleted through 20% sucrose cushions. Cells were collected and lysed in 0.5% lysis buffer (RIPA Buffer) (140 mM NaCl, 8 mM Na2HPO4, 2 mM NaH2PO4, 1% Nonidet P-40 [NP-40], 0.5% sodium deoxycholate, 0.05% sodium dodecyl sulfate [SDS]) and Complete Protease Inhibitor Cocktail (Roche). Isolated virions and viral proteins in cell lysates were analyzed by SDS-polyacrylamide gel electrophoresis (PAGE) and Western blotting for viral proteins with a goat anti-HIV NC (a generous gift from Robert Gorelick) and a mouse anti-CA antibody (AIDS Repository cat# clone 183-H12-5C). Host proteins were detected by Western blotting with mouse anti-tubulin and VPS28 expression was verified with a rabbit anti-VPS28 antibody (Sigma).

### CG Model Generation of ESCRT-I

All-atom (AA) molecular dynamics of the human headpiece of ESCRT-I and UEV domain of TSG101 were performed for the CG model generation of ESCRT-I. Initial AA configurations of the human headpiece of ESCRT-I were generated from the crystal structure reported in this study. Initial AA configuration for the UEV domain of TSG101 was prepared from its crystal structure available in the Protein Data Bank [PDBID: 3OBU].^32^ The systems for AA simulation were prepared using the tleap module in Amber18 MD software.^70^ To generate a solvated protein system, the protein was placed in a pre-equilibrated water box with at least a 1.5 nm layer of water from the surface of protein to the edge of the simulation cell. Counterions were added to compensate for any net charge on the protein resulting in a charge neutral system. The final system of ESCRT-I headpiece and UEV domain contained 89,070 and 44,874 atoms, respectively. In all simulations, the protein was modeled with Amber ff14SB force field^71^ and water was modeled as TIP3P,^72^ using a 2 fs MD timestep. All bonds between a heavy atom and hydrogen in the protein were constrained using the LINCS algorithm. Each solvated protein system was initially minimized using the steepest descent algorithm for a maximum of 50000 steps or until the maximum force was less than 239 kJ/mol/nm. This was followed by a short equilibration of 1 ns in the constant *NVT* ensemble, during which the heavy atoms of the protein were constrained relative to corresponding initial position with a harmonic restraint of 239 kcal/mol/nm^2^. The temperature was maintained at 310 K using stochastic velocity rescaling thermostat with a coupling constant of 2 ps. A further 1 ns of equilibration was performed in the constant *NpT* ensemble to relax the initial simulation cell. The pressure was maintained at 1 bar using the Berendsen barostat with a coupling constant of 10 ps. Production runs of 500 ns for each system were then performed in the *NpT* ensemble by removing all constraints on heavy atoms. In the production runs, temperature was maintained at 310 K using a Nose-Hoover thermostat with a coupling constant of 2 ps; pressure was maintained using Parrinello-Rahman barostat with a coupling constant of 10 ps. The final AA simulations were performed using the Gromacs 2018 MD software package.^73^

The CG model of ESCRT-I headpiece and UEV domain of TSG101 was generated from the corresponding all-atom trajectories using the Essential Dynamics Coarse-graining (ED-CG) method.^74^ ED-CG is a bottom-up method that variationally determines CG sites to capture the fluctuations in the essential subspace determined from principal component analysis of the all-atom trajectories. The resulting CG model of ESCRT-I headpiece and UEV domain of TSG101 have 33 and 19 CG sites respectively, with an average resolution of ∼8 amino acids per CG site. Next, effective interactions between CG sites were represented as a network of effective harmonic spring for all neighboring CG sites within 3 nm of the reference CG site. The spring constants for CG site pairs were derived using the hetero-elastic network (hENM) method.^75^ The CG model for the ESCRT-I stalk is directly derived from the experimental crystal structure for yeast ESCRT-I available in Protein Data Bank [PDBID: 2P22]^35^ maintaining the same average resolution of ∼8 amino acids per CG site. Yeast ESCRT-I stalk contains 3 subunits - Vps23, Vps37 and Mvb12. In the case of 4 long α helices constituting the stalk [Vps23 (α1, α2), Mvb12 (α2) and Vps37 (α4)], a single CG site is created every 8 consecutive amino acids. Two additional CG sites were placed to represent Vps37 (α1, α2) at the base of the stalk. The initial position of CG sites was calculated from the center of mass of the Cα atom of the corresponding amino acid residues that the CG site represent. Overall, the CG model of the stalk contains 31 CG sites. Next, all CG sites within 3 nm of a central CG site were connected with effective harmonic springs. In the case of CG sites corresponding to the same subunit (e.g., Vps23 and Vps23) we used a spring constant of 50 kcal mol^-1^ nm^-2^, while for CG sites corresponding to different subunits a weaker spring constant of 5 kcal mol^-1^ nm^-2^ was used. The stalk and the headpiece were connected by harmonic bonds using the same protocol as above. Finally, inter-domain interactions were represented using a combination of a soft excluded volume (*E_excl_*) and Gaussian attractive (*E_gauss_*) potentials where *r_ij_* is the pairwise distance between CG site types *i* and *j*. For all CG sites types at headpiece involved in self-assembly Gaussian attractive (*E_gauss_*) potentials were used. Using the CG-mapped structure of the 12-mer ESCRT-I headpiece, the mean distance between a pair *ij* is assigned as *r_ij,h_*. The *r_ij,h_* values ranged from 1.28 to 1.8 nm. For these attractive *ij* pairs *σ_ij_* = 0.09 nm, and *H_ij_* = −0.9 nm kcal/mol are used. We note that strength of ESCRT-I attractive interaction (*E*_E1-E1_) described in the main text is same as 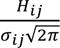. The particular chosen value of *H_ij_* is the maximum value that does not lead to significant oligomerization in ESCRT-I solution in the CGMD simulations (Fig. 6b). This leads to the *E*_E1-E1_ value of −4 kJ/mol. Details of the simulations of ESCRT-I solutions are described in the section having the CG simulation details. All pairwise interaction modeled with *E_gauss_* used a 30 Å radial cutoff. For the rest of the pairwise CG interactions a soft excluded volume (*E_excl_*) interaction is used with *A_ij_* = 25 kcal/mol, *r_ij,c_* = 1.5 nm. The linker domain is represented as a linear chain of 9 CG particles that are linked with harmonic bonds of spring constant 50 kcal mol^-1^ nm^-2^ and equilibrium distance is maintained at 1.5 nm. The intra- or inter-domain interaction of the linker CG sites is modeled with excluded volume (*E_excl_*) potential with the same parameters as above. To model the binding of the p6 domain of Gag and UEV domain of TSG101, we first added an extra CG binding site at the end of the NC domain of the CG Gag (the details of the CG Gag model is described in the next section). This CG binding site is linked to three nearest CG beads of NC domain by harmonic bonds of spring constant 50 kcal mol^-1^ nm^-2^ and equilibrium distance 1.5 nm. To implicitly allow binding of ESCRT-I and p6 domain, 2 C-terminal CG sites of UEV domain of TSG101 with the binding site. The attractive interaction is modeled with Gaussian attractive (*E_gauss_*) potentials, with values *r_ij,h_* = 1.2nm, *σ_ij_* = 0.09 nm, and *H_ij_* = −4.2 nm kcal/mol are used. In a separate publication details of the characterization of the ESCRT-I CG model will be presented.

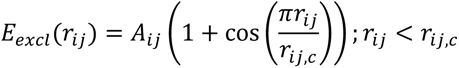

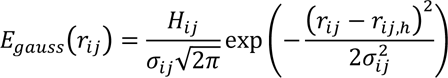

### CG Model Generation of Gag

Prior to CG model generation, AA molecular dynamics simulations of the matrix (MA), capsid/spacer peptide 1 (CA/SP1), and nucleocapsid (NC) domains were performed using either Amber18 (for MA and NC)^70^ or Gromacs 2016 (for CA/SP1).^73^ Initial protein configurations were adopted from atomic models for MA (PDB 2H3I),^76^ CA/SP1 (PDB 5L93),^77^ and NC (PDB 5I1R);^78^ in the case of CA/SP1, an 18-mer configuration in the form of a hexamer of trimer of dimers was used. All proteins were solvated by water and 150 mM NaCl in a cubic simulation domain large enough to contain a 2.5 nm layer of water beyond the protein-solvent interface. Energy minimization is performed using steepest descent until the maximum force is less than 239 kcal/mol/nm. Equilibration is performed with harmonic restraints (using a 239 kcal/mol/nm^2^ spring constant) on each heavy atom throughout the protein for 1 ns in the constant *NVT* ensemble using a Langevin thermostat for MA and NC (stochastic velocity rescaling thermostat for CA/SP1) at 310 K and a damping time of 1 ps. Restraints were then removed and each system was allowed to equilibrate for 100 ns in the constant *NPT* ensemble using a Nosé-Hoover chain thermostat with a 2 ps damping time and a Monte Carlo barostat for MA and NC (Parrinello-Rahman barostat for CA/SP1) with a 10 ps damping time at 310 K and 1 bar. Production runs were performed over 2.4 µs for MA, 2.2 µs for NC, and 0.9 µs for CA/SP1 with configurations saved every 100 ps. All simulations used the CHARMM36m^79^ and TIP3P^72^ force fields for protein and water, respectively, with an AA MD timestep of 2 fs. Hydrogen containing bonds were constrained using SHAKE for MA and NC and LINCS for CA/SP1.

The CG model for Gag used in this work is meant to emulate the morphology of an immature lattice. To this end, each of the three protein domains are coarse-grained from AA molecular dynamics trajectories using the following procedure. First, each protein sequence is partitioned into *N* CG sites (*N* = 20, 35, and 11 for MA, CA/SP1, and NC, respectively) using the ED-CG method.^74^ Next, intra-domain interactions are represented as a heterogeneous network of effective harmonic springs for all CG pairs within a cutoff of 1.6 nm for MA and NC and 1.8 nm for CA/SP1; here, spring constants were iteratively optimized to minimize the difference between the normal mode fluctuations of the network and the fluctuations from the AA MD trajectories. The MA-CA/SP1 and CA/SP1-NC domains are connected by bead-spring chains of length 9 and 2, respectively, with a spring constant of 50.0 kcal/mol/nm^2^ and equilibrium distance of 0.7 nm. Finally, inter-domain interactions are represented using a combination of a soft excluded volume (*E_excl_*) and Gaussian attractive (*E_gauss_*) potentials. For the majority of the pairwise CG interactions, we used *A_ij_* = 25.0 kcal/mol, *r_ij,c_* = 0.9 nm, *H_ij_* = 0 nm kcal/mol, *σ_ij_* = 0 nm, and *r_ij,h_* = 0 nm. The latter four parameters are customized for pair interactions involving the CA/SP1 domain, which is known to be the domain associated with protein-protein associated. Using the CG-mapped trajectories of the CA/SP1 18-mer, statistics on the inter-protein distances (for *r_ij_* < 2 nm) between each pair of CG site types are gathered. For each *i* and *j*, the minimum distance are used for *r_ij,c_*. Gaussian functions (= *C*exp[-(*r*-*r_h_*)^2^/(2*σ*^2^)]) are fit to each distribution of *r_ij_*_;_ the subset of CG pairs with *r_h_* < 1.5 nm and *σ* < 0.15 nm were considered attractive interactions, i.e., close-contact pairs with minimal fluctuations, such that *r_ij,h_* = *r_h_*, *σ_ij_* = *σ*, and *H_ij_* = −0.6 nm kcal/mol. These interaction parameters are found to be sufficient to stabilize the formation of the immature lattice, which we tested by aligning hexamers of our CG model to cryo-electron microscopy maps of immature HIV-1 lattices provided by John Briggs (MRC LMB). Full details and characterization of the CG Gag model will be presented in a future publication.

### CG Simulation details and analysis

All CGMD simulations were performed using the LAMMPS MD software^80^. The equations of motions were integrated with the velocity Verlet algorithm using a time step (*τ*) of 50 fs. The simulations of ESCRT-I assembly templated by Gag shell were performed under constant *Np_xy_T* ensemble. Periodic boundary conditions were implemented in the direction parallel to the membrane plane (*xy* plane). The simulation cell was nonperiodic in the *z* direction and terminated by a reflective wall. In the simulation setup with the “budding neck” geometry, the membrane region in contact the Gag shell were kept frozen, while the equations of motion for the rest of the membrane patch was integrated. This allows to maintain the “bud-like” shape of the membrane. Similarly, for the CGMD simulations for ESCRT-I self-assembly, the MA-CA/SP1 domain of each Gag polyprotein is kept frozen to maintain the crystallinity of the relaxed Gag shells. The temperature in the simulations was maintained at 310 K using Langevin dynamics with a damping constant of 200*τ* ^81^. The pressure in the *xy* direction was controlled with the Nose-Hoover barostat with a coupling constant of 500*τ* ^82^. The pressure tensors in the *x* and *y* directions were coupled and maintained at 0 bar. The simulations of ESCRT-I in solution were performed in the constant *NVT* ensemble. The system temperature in these simulations was maintained in the same way as described above, while the Gag and ESCRT-I were modelled with the CG models described earlier. The lipid bilayer was modeled with a 3-site CG model ^83^. Visualization of the simulation trajectory was created using Visual Molecular Dynamics (VMD) software ^84^. We characterized ESCRT-I oligomerization by applying a neighboring criteria: an ESCRT-I hetero-tetramer (*i*) was considered to be bound to another hetero-tetramer (*j*) if all the binding CG pairs *ij* at the headpiece were within a 0.3 nm cutoff.

We prepared model Gag shells used in the simulation of ESCRT-I assembly from cryo-electron microscopy maps of immature HIV-1 lattices, with an approximately hemispherical geometry. These immature Gag lattices were mapped to CG representation with our CG Gag model and relaxed. Using the relaxed structure as our parent lattice, we selected a segment of the hemisphere, typically of height 20 nm. The upper and lower limit of the height used to select a segment was progressively varied to create Gag shells of different diameters (*D*_Gag_). We note that *D*_Gag_ in this study was defined as the lowest cross-section of the segment. The Gag shells were then placed in a simulation cell of dimensions 152 nm × 152 nm × 56 nm in the *x*, *y*, *z* directions respectively, as depicted in Fig. 6c. Then, the membrane layer resembling a “bud-like” shape was wrapped around the Gag shells. The composite systems were then relaxed for 5 × 10^6^ CGMD timesteps. We performed three independent simulations of ESCRT-I self-assembly for each Gag shell (4 different cross-sections) resulting in 12 total simulations. In each simulation the initial velocity distribution of inserted ESCRT-I was assigned via a random number generator seed from Boltzmann distribution. Simulation trajectories were saved every 1 × 10^6^ CGMD timesteps.

To prepare systems of ESCRT-I solution we first created a simulation cell of dimension 200 nm × 200 nm × 200 nm. Next in the simulation cell ESCRT-I units are evenly spaced in a 3-dimensional grid of (4 × 4 × 4) or (4 × 4 × 8) with a 10 nm separation between every ESCRT-I unit, resulting in two systems containing 32 or 64 ESCRT-I heterotetramers. Then 3 independent simulations were evolved at 310 K for each system (32 or 64 ESCRT-I) for 10 × 10^6^ CGMD timesteps. In these simulations all attractive interactions between ESCRT-I headpiece is turned off to prevent oligomerization. The final state of each independent system is used as initial configuration for simulations of ESCRT-I assembly in solution. For the assembly simulations, we turn on the attractive interactions between ESCRT-I headpiece and vary the value of *E*_E1-E1_ from −3.1 to −4.9 kcal/mol (*H*_ij_ from −0.7 to −1.1 nm kcal/mol). For each *E*_E1-E1_, three independent simulations are evolved for 200 × 10^6^ *τ* resulting in a total of 15 simulations.

## Accession codes

Coordinates and structure factors have been deposited in the Protein Data Bank under accession code PDB 6VME.

## Acknowledgments

We thank B. Yang for contributing to early stages of this project, S. Fromm, C. Buffalo, and P. Grob for electron microscopy advice and support, K. Larsen for comments on the manuscript, and J. Briggs for providing immature Gag lattice maps. This work was supported by NIH grants R37 AI112442 (J.H.H.), R01 GM127954 (H.G.W), R01 GM128507 (A.H. and G.A.V.) and F32 AI150477 (A.J.P.). Beamline 8.3.1 at the Advanced Light Source is supported by the National Institutes of Health (R01 GM124149 and P30 GM124169), Plexxikon Inc., and the Integrated Diffraction Analysis Technologies program of the US Department of Energy Office of <u>Biological and Environmental Research</u>. The Advanced Light Source (Berkeley, CA) is a national user facility operated by Lawrence Berkeley National Laboratory on behalf of the US Department of Energy under contract number DE-AC02-05CH11231, Office of Basic Energy Sciences.

## Author contributions

Conceptualization, T.F., Y.T., G.V., J.H.H.; Investigation, T.F., Y.T., A.H., K. R., N.T., A.L.Y., X.L.; Resources, A.P.; Supervision, H.G.W., F. B., G.V., J.H.H.; Writing-original draft, T.F., Y.T., A.H., G.V., J.H.H.; Writing-review and editing, all authors.

---

**Movie S1: CGMD simulation of ESCRT-I assembly** Assembly of ESCRT-I templated by a Gag shell with a 54 nm opening. The color representation is same as described in Fig. 6. For each frame only the largest ESCRT-I oligomer is shown. We note that for some frames there are two largest oligomers of same size, in that case both the oligomers are shown. The membrane is not rendered for visual clarity.

**Fig. S1:**
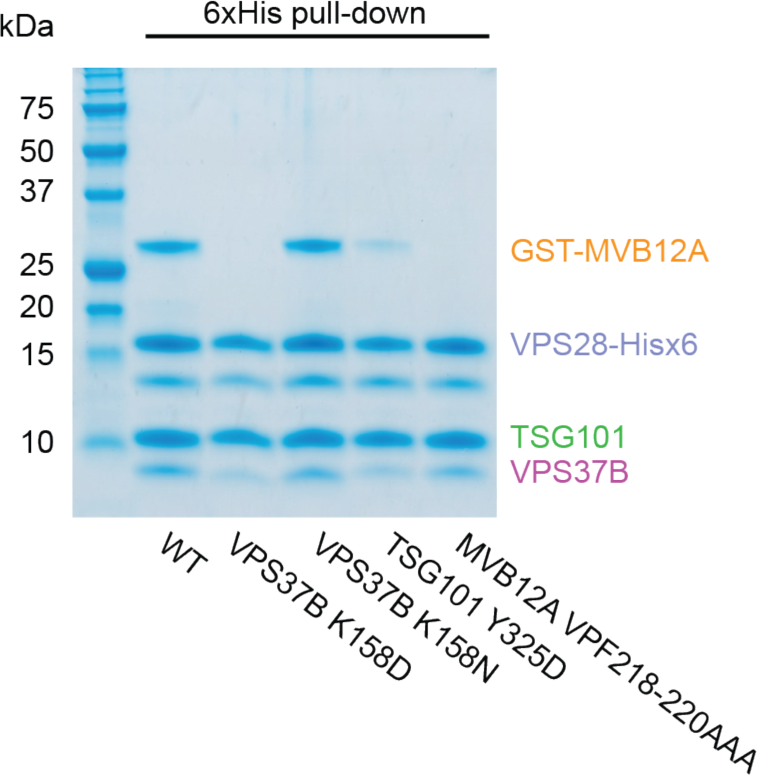
Effect of specific mutations on ESCRT-I head complex integrity Mutant versions of the ESCRT-I head complex were expressed in *E. coli.* VPS28 and MVB12A subunits were expressed as C-terminal hexahistidine and N-terminal GST fusions respectively. Lysate was incubated with Ni-NTA agarose beads and complex integrity analyzed by SDS-PAGE following multiple washes.

**Fig. S2:**
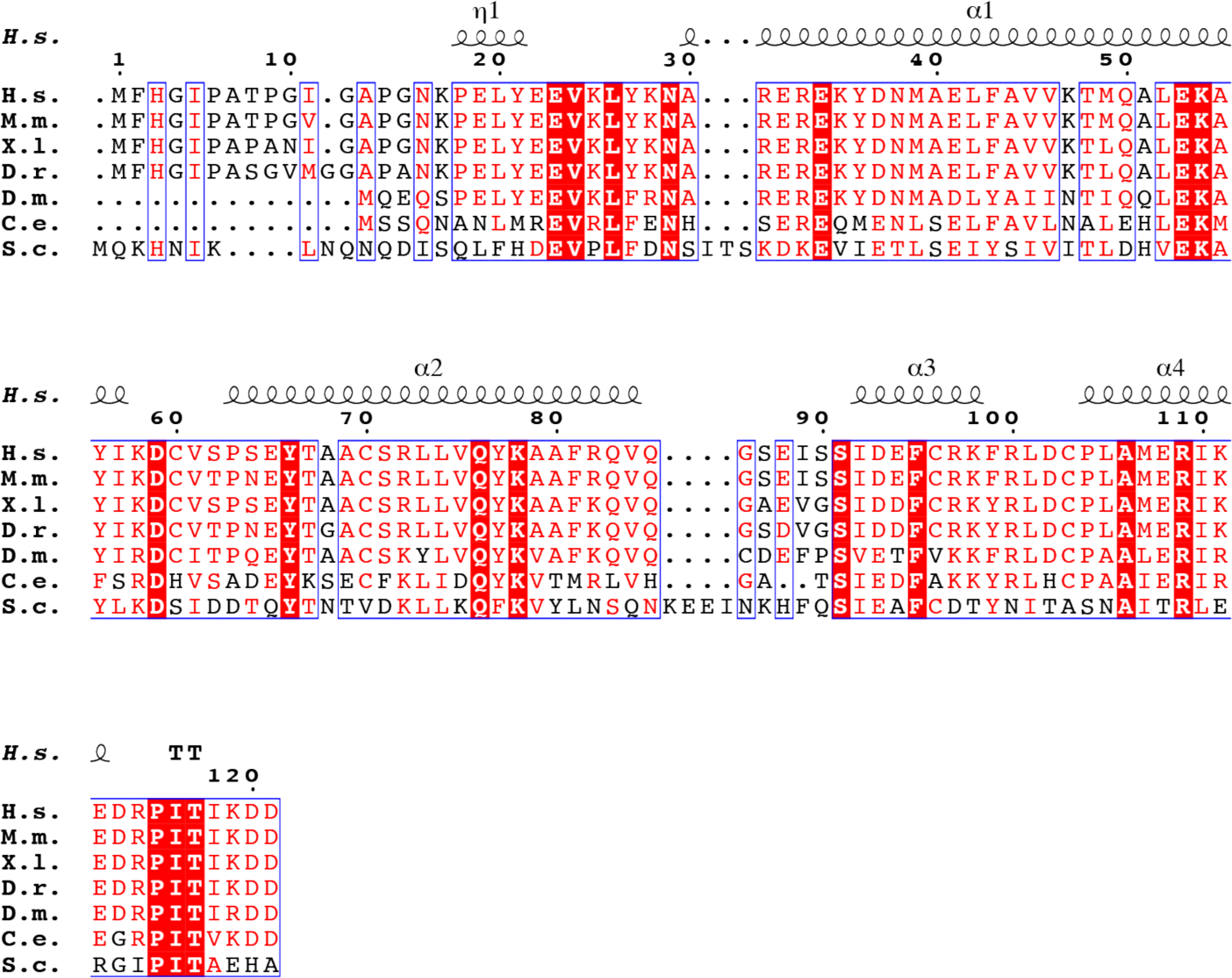
Sequence alignment of VPS28 orthologs Secondary structure displayed above the alignment is derived from the human ESCRT-I head structure. Alignment was generated using ClustalW and ESPript.

**Fig. S3:**
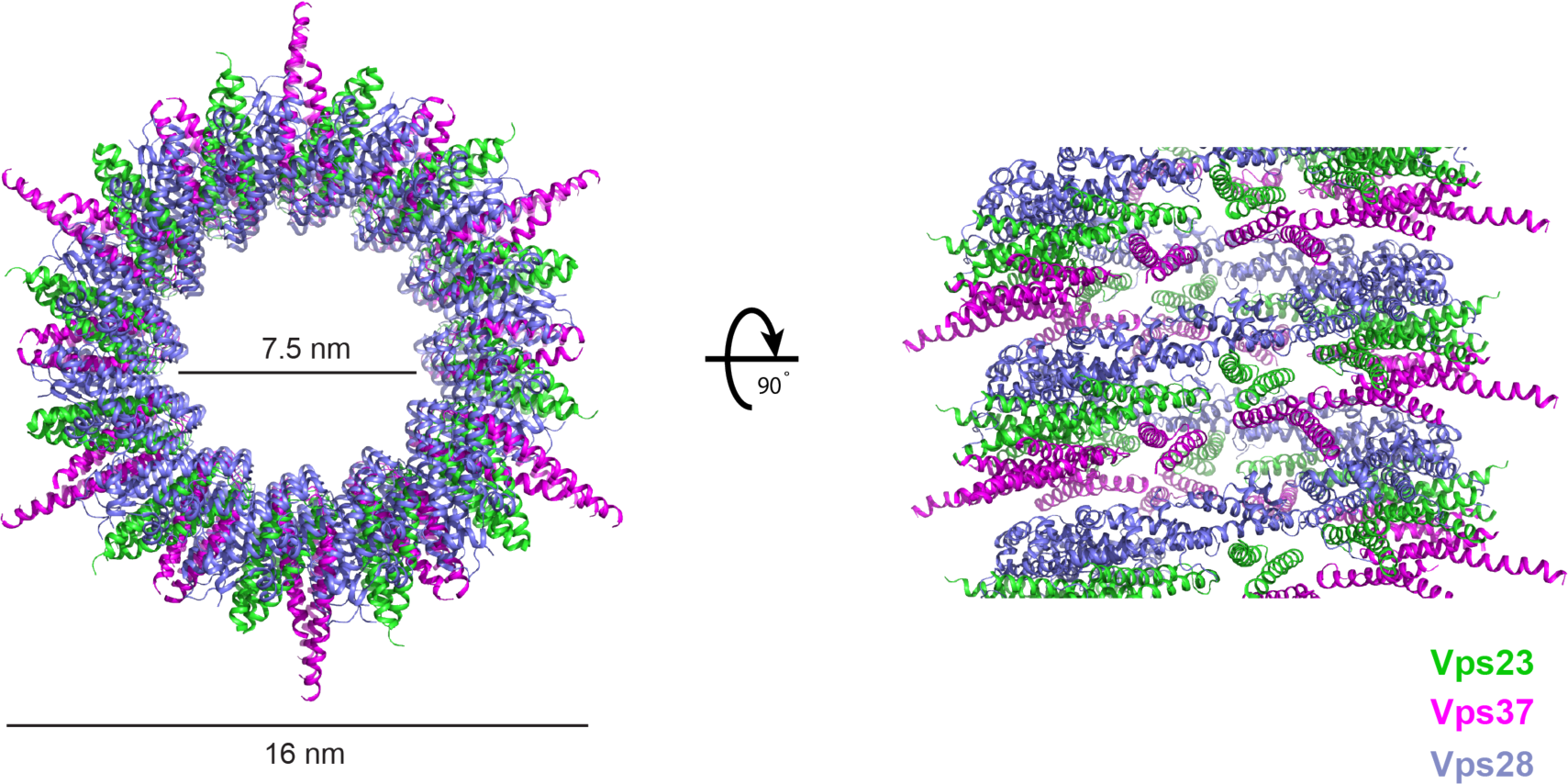
Helical yeast ESCRT-I head tubes The trimeric yeast ESCRT-I head forms helical tubes within a crystal (PDBID: 2CAZ). The crystal is constructed from a series of laterally stacked tubes where each tube is composed of a single, continuous helix of yeast ESCRT-I head protomers. Vps23, Vps28, Vps37 are colored green, purple and magenta respectively. Tube dimensions are labeled.

**Supplementary Table 1.**
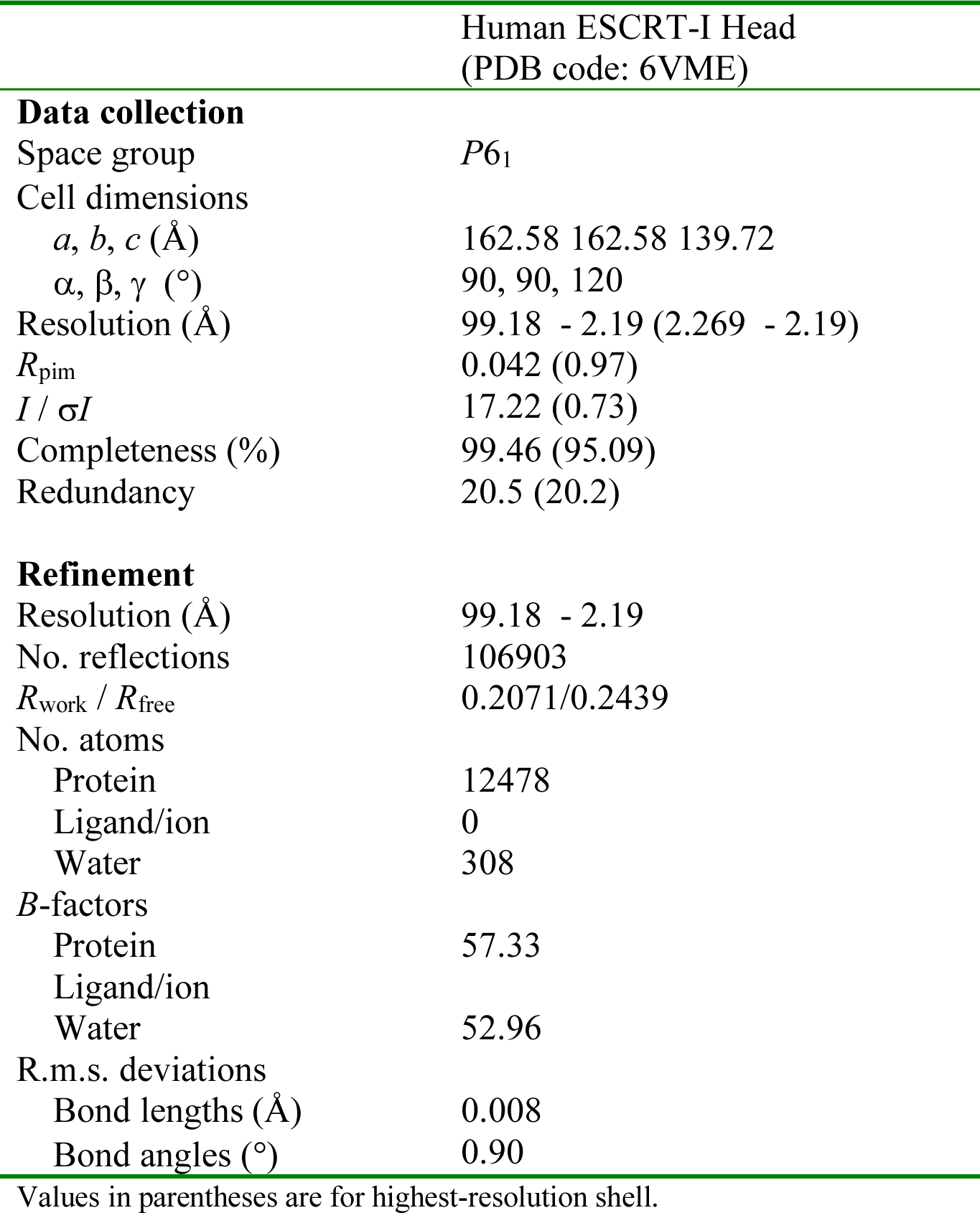
Data collection and refinement statistics

